# Carcass Scavenging Relaxes Chemical-Driven Female Interference Competition

**DOI:** 10.1101/2020.06.02.130146

**Authors:** Basabi Bagchi, Srijan Seal, Manasven Raina, Dipendra Nath Basu, Imroze Khan

**Affiliations:** Ashoka University, Plot No. 2, Rajiv Gandhi Education City, National Capital Region, P.O. Rai, Sonepat, Haryana-131029, India; National Centre for Biological Sciences, UAS-GKVK, Bellary Road, Bangalore 560065, India

**Keywords:** Carcass-scavenging, Carcass-quality, Chemical interference, Female density, Intra-sexual competition, Protein access, Quinones

## Abstract

Female-female nonsexual interference competition is rapidly emerging as a major fitness determinant of biased sex-ratio groups with high female density. How do females overcome such competition? We used adult flour beetle *Tribolium castaneum* to answer this question, where females from female-biased groups suppressed each other’s fecundity by secreting toxic quinones from their stink glands, revealing a chemical-driven interference competition. The added natal resource did not alleviate these fitness costs. Females also did not disperse more at high female-density. Hence, the competition was neither limited by the total resource availability nor the inability to avoid chemical interference. Instead, protein sequestered via scavenging of nutrient-rich carcasses relaxed the female competition, by increasing their fecundity and reducing the quinone content. Even infected carcasses were scavenged to extract fitness benefits, despite the infection-risk. Finally, individual stink gland components triggered carcass-scavenging to increase fecundity, indicating a potentially novel chemical feedback loop to reduce the competition.

## INTRODUCTION

In many animals, females compete strongly for resources such as food or space, rather than mates, using various interference strategies (reviewed in Stockley & Bro-Jørgensen 2011; Tobias *et al.* 2012). For instance, high-ranking female chimpanzees (Murray *et al.* 2006) or meerkats (Bell *et al.* 2014) monopolise access to high-quality diet by dominance relationships or territorial aggressions, thereby suppressing the reproduction of subordinate females to maximise their own reproduction. In contrast, females parasitoids use biochemical mechanisms while competing for a suitable resource (Harvey *et al.* 2013). These multifaceted competitive interactions are further exemplified by recent experiments with flour beetles where females compete using defensive quinones as the secreted toxin in response to female density rather than via direct intra-sexual interactions (Khan *et al.* 2018). Females inhibit each other’s reproduction, either due to their own secreted chemicals or due to chemicals secreted by other competing females.

How do females overcome fitness effects of interference competition? As food availability is a limiting factor for female fitness in many species (Ho & Dawson 1966; Wheeler 1996), one possibility is that fitness impacts of interference competition can be directly reversed by increasing the access to the available natal resource. Alternatively, animals can also disperse from competitive environments (Waser 1985; Porter & Dooley 1993; Matthysen 2005), using information acquired from their external surroundings (Clobert *et al.* 2009; Clutton-Brock & Lukas 2012). While the relative importance of dispersal during density-dependent female interference competition is still unclear, environmental cues affecting dispersal decision are widely known. For instance, mice trigger their dispersal, using the odour of urine, which serves as a cue for relatedness and competitiveness of neighbouring conspecifics (Isles *et al.* 2002; Latham & Mason 2004). In *Tribolium* beetles, dispersal rate increases in response to the conditioning of flour by the accumulation of excreta or secreted quinones at high population density (Ogden 1969). Dispersal from high-density patches can increase further if beetles are housed with individuals that are reared under poor conditions as juveniles (Endriss *et al.* 2019), suggesting that dispersal behaviours can even be sensitive to the environmental history of neighbouring conspecifics. These results might also have direct implications for female dispersal during competition. For instance, density-dependent accumulation of defensive chemical secretions, as observed in flour beetles (Khan *et al.* 2018), can stimulate female dispersal to avoid the overall negative fitness consequences of female-biased sex-ratios.

Finally, facultative scavenging of animal carcasses can be another ecologically relevant and low-cost (with no efforts for predation) foraging strategy (Devault *et al.* 2003; Moleón *et al.* 2014) to reduce competitive interactions by sequestering additional energy and nutrients. Decomposing flesh of protein-rich carcasses (Huntington & Higley 2010) can provide easy access to essential nutrients to stimulate reproductive potential (Piper *et al.* 2014; Corrales-Carvajal *et al.* 2016). Due to these inherently low costs of acquisition and high nutritional value of carcasses, scavenging is perhaps more effective to avoid female competition than access to the natal resource or costly dispersal strategy (Li & Kokko 2019). However, despite the importance of carcass scavenging as an integral part of the energy and nutrient flow model (Wilson & Wolkovich 2011; Turner *et al.* 2016; Abernathy *et al.* 2017), its direct role in female competitive interferences has been rarely analysed (but see Pusceddu *et al.* 2018). Notably, we have no information on whether and how scavenging can influence unique chemical interference competitions observed in some insect species (Harvey *et al.* 2013; Khan *et al.* 2018). Since chemicals act as indicators of female competition in these insects, it is possible that they can also signal increased scavenging to acquire more nutrients and reduce the competition, using a chemical feedback loop.

It is also important to note that a significant fraction of available carcasses in nature is derived from animal deaths due to infection and disease (Devault *et al.* 2003). Although these infected carcasses still serve as an essential nutrient source, overall quality might be poor due to excessive microbial growth (Milutinović *et al.* 2015). However, these carcasses might not only reduce the net benefits of scavenging, but they can also increase the risk of disease transmission in populations (Borchering *et al.* 2017; Sage *et al.* 2019). To counter these effects, organisms might show strong avoidance to scavenging of infected carcasses. However, it is unclear whether, or to what extent, such variation in carcass-quality and scavenging behaviour affect fitness during competition. In the present work, we tested multiple novel hypotheses vis-à-vis analysing the relative impacts of diverse counter-strategies such as natal resource access, increased dispersal, and carcass scavenging to reduce the density-dependent female interference competition. We used *T. castaneum* since they are already an established model system to study female competition (Khan *et al.* 2018; Halliday *et al.* 2019). Wild populations of *Tribolium* beetles are also generalised feeders, show cannibalism (Via 1999) and can survive as natural predator and scavengers of other insects in a range of habitats such as nests of birds or eusocial insects and stored product environments (reviewed in Dawson 1976; Alabi *et al.* 2008). Finally, detailed information on their stink gland secretion (Li *et al.* 2013) and fitness effects (Khan *et al.* 2018) further make them an attractive model to test chemical regulation of scavenging or dispersal behaviour directly. We asked three major questions: I. Is protein-rich carcass scavenging the most effective strategy to avoid adverse fitness effects of female competition? II. Can carcass-quality determine the fitness benefits of scavenging? III. Finally, can chemicals involved in the interference competition regulate scavenging behaviour to increase fitness effects? Overall, our results identified carcass scavenging as the most effective strategy to relax the female competition, with new implications for the evolution of intra-sexual resource competition, chemical ecology and spread of disease in the wild.

## MATERIALS AND METHODS

We used an outbred laboratory population of red flour beetle *Tribolium castaneum* to generate the experimental beetles (See Khan *et al.* 2018 for detailed maintenance protocol). In most experiments, we manipulated female density by grouping four virgin adults (10-day-old) into the following sex-ratio treatments, unless stated otherwise: (1) Male-biased groups (MB): 3 males and 1 female and (2) Female-biased groups (FB): 1 male and 3 females. We housed each replicate group in a 35mm Petri-plate with 3gm of wheat flour to minimise resource limitation. After 8 days, we used them for various experiments as described below (see Fig. 1 for a brief experimental outline). We have provided the details of data analyses in the supplementary information (SI).

**Fig 1.**
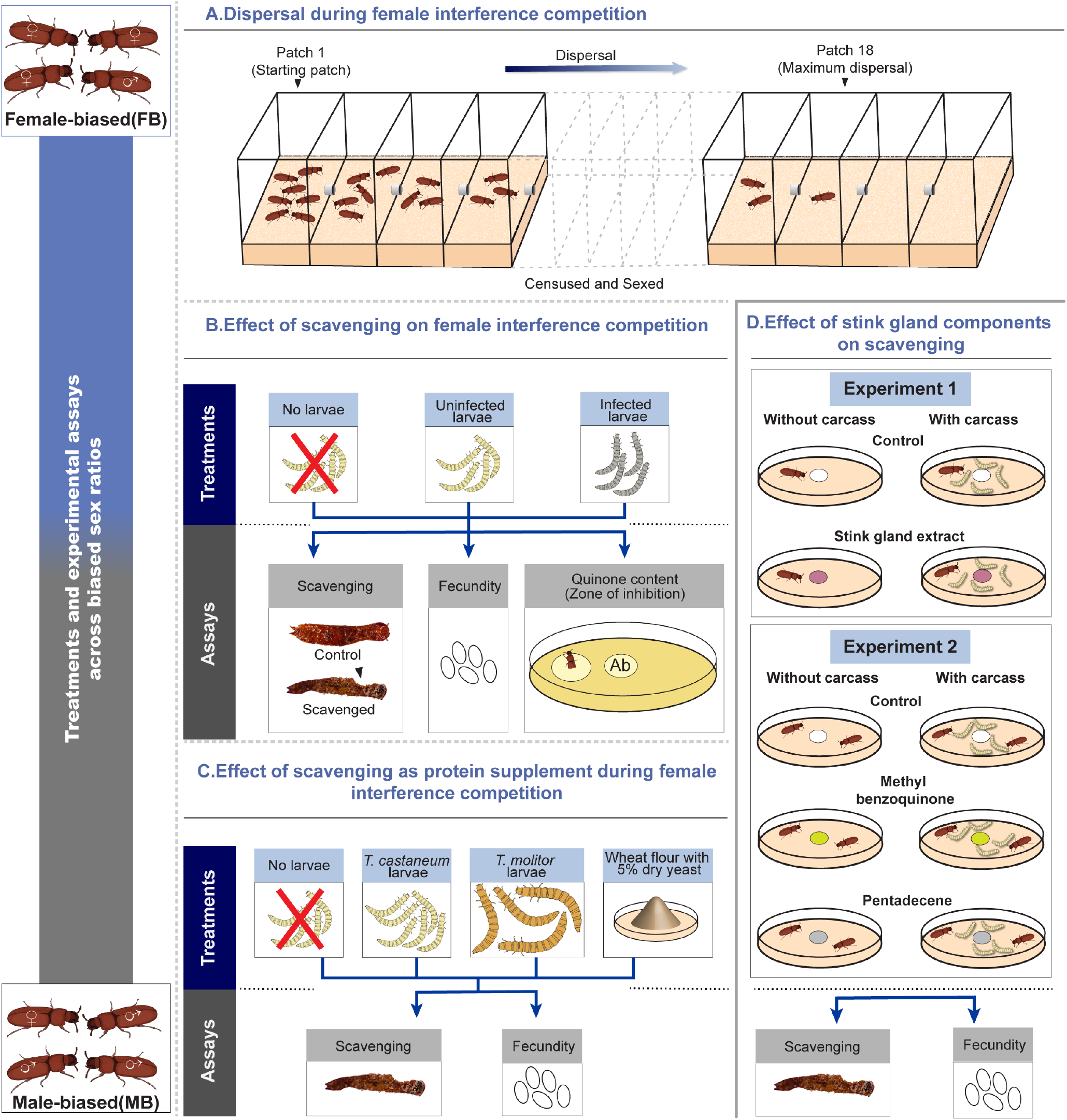
A brief outline of the experiments. (**A) Dispersal during female interference competition**. Male-biased (10 females + 30 males) and female-biased (30 females + 10 males) sex-ratio groups were constituted and added to the first patch of the linear arrays (connected with 2mm holes, described in Endriss *et al.* 2019) to disperse for 30 hours. Following this, beetles were sexed and censused within each patch to estimate sex-specific dispersal from the patch of origin (i.e. patch 1) (n=10 replicates/sex-ratio bias). (**B) Effect of scavenging on female interference competition.** Male-biased (1 female + 3 males) and female-biased (3 females + 1 male) sex-ratio groups were constituted, with or without the access to normal or infected *T. castaneum* larval carcasses. After 8 days, females were assayed for scavenging activity (measured as the fraction of larval carcass remaining) (n=9-10 replicates/sex-ratio bias/carcass type), fecundity (n=9-10 replicates/sex-ratio bias/carcass type/replicate experiment) and quinone production (quantified as zone of inhibition (ZIs) produced by quinones released from females after the cold-shock) (n=12-19 replicates/sex-ratio bias/carcass type). ZIs produced by each female was normalised by dividing with ZIs produced by antibiotic Streptomycin added at the centre of the respective Petri-plates to minimise the variations across plates during experimental handling. (**C) Comparing the effects of scavenging and dry yeast as a protein supplement during female interference competition.** Different sex-ratio groups (constituted as described above) received carcasses derived from *T. castaneum* or *T. molitor* larvae or, 5% active dry yeast and 8 days later, they were assayed for female scavenging activity and fecundity (19-20 replicates/protein source/sex-ratio bias/assay). (D**) Effect of total stink gland extract and individual components on scavenging activity and fecundity.** 10-day-old 2-4 mated test females (reared under 1:1 sex-ratio), with or without the access to *T. castaneum* larval carcasses, were exposed to paper discs soaked with total stink gland extract (treatment 1) or its components such as methyl-p-benzoquinone or 1-pentadecene (treatment 2) for 8 days. Subsequently, they were assayed for scavenging activity and fecundity (n=18-20 groups/stink gland extract or components/assay).

### Estimating the effects of natal resource on female competition

We directly quantified the fitness effects of total resource during interference competition by varying the quantity of wheat flour available per adult. We constituted MB and FB groups using 10-day-old virgins as described above and provided groups with 1g, 4g, or 8g flour per beetle (*n* = 10 replicates/resource-level/sex-ratio). We measured female fecundity after 8 days, where one female from each sex-ratio group was allowed to oviposit in 5gm of sifted wheat flour in a 60mm Petri-plate. After 24 hours, we removed the females and sifted the flour using a 300μm sieve (TIC Test Sieves) to count the number of eggs.

### Quantification of dispersal under biased sex-ratios

Flour beetles possess paired abdominal and thoracic stink glands that contain chemicals such as benzoquinones (e.g. methyl-(MBQ) and ethyl-benzoquinone (EBQ)) and hydrocarbons (e.g. 1-pentadecene (PD)) (Li *et al.* 2013). Our previous data suggested that increased production of benzoquinones can directly mediate female interference competition in FB groups (Khan *et al.* 2018). We thus directly compared the dispersal behaviour of FB vs MB beetles to test whether FB females dispersed more to avoid such chemical-driven interference competition. We closely followed a protocol developed by Endriss *et al.* (2019), measuring beetle dispersal in linear arrays of twenty 4 × 4 × 6 cm replicate plastic boxes (or patches) that were connected by 2mm holes (See **Fig.** 1A). We have provided a detailed protocol in the SI. We constituted beetle groups with similar sex-ratios as described above, but with a larger group size of 40 individuals to obtain a meaningful comparison of dispersal probability across sex-ratios groups (Endriss et al. 2019). We added beetles to the first patch of linear arrays (n = 10 replicates/sex-ratio). After allowing them to disperse for 30 hours, we counted all experimental males and females within each patch across all the arrays to estimate how many patches they dispersed from the patch of origin.

Separately, we also tested the causal link between quinones and dispersal by analysing the direct effect of MBQ on the dispersal of beetle groups comprised of 10 males and 10 females (n=9 replicates/treatment) (See SI methods).

### Estimating the effects of scavenging on female competition

#### Generating larval carcasses

To generate carcasses, we mostly sacrificed 14-day-old *T. castaneum* larvae by freezing them at −20°C for 30 minutes. Since cannibalism of juveniles is widely known in *Tribolium* adults (Sonleitner & GuthJanis 1991; Via 1999), we expected that availability of conspecific larval carcasses would ensure a high consumption rate. We also used 20-day-old *Tenebrio molitor* larvae, a closely related species to *T. castaneum* (Sokoloff 1977) to verify that the observed fitness effects are not only limited to cannibalism (discussed below).

We manipulated the carcass quality by mimicking animal death due to infection, where 14-day-old *Tribolium* larvae were first infected with a lethal dose of natural beetle pathogen *Bacillus thuringiensis* (Bt) (Abdel-Razek *et al.* 1999). Detailed infection protocol has been described previously (Khan *et al.* 2017). Since most larvae died within the first 8-10 hours of infection due to rapid Bt growth, we collected larvae that died within 6 hours of infection. Separately, we also quantified the beetle avoidance to these infected carcasses, using a two-way choice experiment where they were simultaneously provided access to both infected vs normal carcasses (See the SI methods, **Fig.** S1).

#### Scavenging activity and fecundity assay

We constituted different sex-ratio groups, with or without the opportunity to scavenge infected vs uninfected larval carcasses (n= 9-10 replicates/sex-ratio bias/carcass type) (**Fig** 1B). We weighed freshly prepared carcasses in groups of 10 larvae (Mean ± SE: 20.18 ± 0.07 mg) and added them immediately to each replicate of sex-ratio groups. After 8 days, we carefully sieved out the remaining body parts of larvae and weighed them again. We calculated the ratio of body weight after and before the exposure to sex-ratio groups, as a proxy for group scavenging where lower ratio (i.e. higher weight loss in the presence of beetles) indicated more scavenging (also see SI for per capita scavenging). We note that a significant amount of weight reduction can also be attributed to the progressive loss of water from carcasses during the long experimental window of 8 days. To quantify this effect of drying alone, we thus included control treatments where carcasses were stored without beetles (i.e. no scavenging) for 8 days. Separately, we also isolated females from each of these scavenging and sex-ratio treatments after 8 days and determined the impacts on their fecundity as described above in two independently replicated experiments (n= 9-10 replicates/sex-ratio bias/carcass-type/replicate experiments).

#### Quinone assay

To estimate the changes in stink gland quinones as a function of both group sex-ratio and scavenging, we set up MB and FB groups as described above (n= 12-19 replicates/sex-ratio/scavenging treatment) (**Fig.** 1B). After 8 days, we estimated stink gland production of one female from each group, relying on the antimicrobial properties of quinones as described in earlier studies (Unruh *et al.* 1998; Khan *et al.* 2015) (also see SI methods). Briefly, we measured the diameter of the zone of inhibition (ZI) produced by individual cold-shocked females embedded vertically in a lawn of Bt grown on agar plates. The cold shock triggered the complete release of the stink gland contents, producing ZIs on agar plates. Our previous data suggested that ZIs are reliable indicators of stink gland quinone contents as they strongly correlate with the methyl- and ethyl-benzoquinone contents of stink glands, quantified by HPLC methods (Khan *et al.* 2018) (see **Fig.** S2).

#### Analysing the importance of protein supplement during interference competition

We next tested whether fitness effects of female competition can be rescued by rich protein sources such as active dry yeast supplement which is also known to increase fecundity in several insects (Perez-Staples *et al.* 2007; Matzkin *et al.* 2011; Michalczyk *et al.* 2011) (**Fig.** 1C). This comparison enabled us to directly correlate the overall fitness benefits of scavenging with the rich protein content of carcasses. Simultaneously, we also tested whether fitness benefits of scavenging are only limited to feeding conspecific larval carcasses (i.e. a form of cannibalistic scavenging; Mckillup & Mckillup 1996), or it can also be extended to other animal carcasses as well. To this end, we first isolated beetles into different sex-ratio groups as described above. We then randomly divided them into the following four treatments groups (n=19-20 replicates/treatment/sex-ratio bias), receiving (1) carcasses of 10 *Tribolium castaneum* larvae (Mean ± SE: 21.72 ± 0.33 mg); (2) 8 *Tenebrio molitor* larvae (Mean ± SE: 21.97 ± 0.51 mg); (3) dietary supplement of 5% yeast in flour; (4) only flour, without the scavenging opportunity or yeast supplementation (control group). We quantified fecundity and scavenging after 8 days as outlined above.

### Quantifying the impacts of stink gland components on scavenging and its fitness effects

Since stink gland components can directly mediate female interference competition (Khan *et al.* 2018), we next tested whether stink gland contents themselves could serve as a trigger for increased scavenging to enhance fitness, using a protocol described in Khan *et al.* (2018) (**Fig.** 1D; see SI methods). In short, we paired two 10-day-old mated standard test females (reared without sex-ratio bias) in a 35mm Petri-plate and exposed them to whole abdominal stink glands extracted in hexane solvent for 8 days and then correlated the changes in scavenging activity with their fecundity (*n*=18-19 females/treatment/scavenging).

Since MBQ and PD are two major benzoquinone and hydrocarbon components respectively of *T. castaneum* stink glands (Unruh *et al.* 1998; Li *et al.* 2013), we next tested whether these individual components are responsible for triggering scavenging behaviour. We thus directly quantified the impact of commercially available MBQ (Sigma) and PD (Sigma) on scavenging and fitness effects, as described above for gland extracts. Briefly, we exposed the females, with or without the access to the carcass, to a filter paper disc (diameter:10mm) soaked with 15μg MBQ or 5μg PD dissolved in 10μl hexane, which was within the physiological range of MBQ and PD content of a single *Tribolium* female (Li *et al.* 2013) and then quantified scavenging and fecundity after 8 days.

## RESULTS

### Natal resource availability did not relax the female competition

We tested the impact of natal resource availability on the female competition by increasing the amount of available wheat flour per beetle from 1gm to 8gm. Although fecundity generally increased with higher resource availability, FB females consistently produced a lower number of eggs compared to MB females, regardless of the resource amount (**Fig.** 2A, **Table** S1A). This suggested a simple increase in natal resource was insufficient to relax the interference competition.

**Fig 2.**
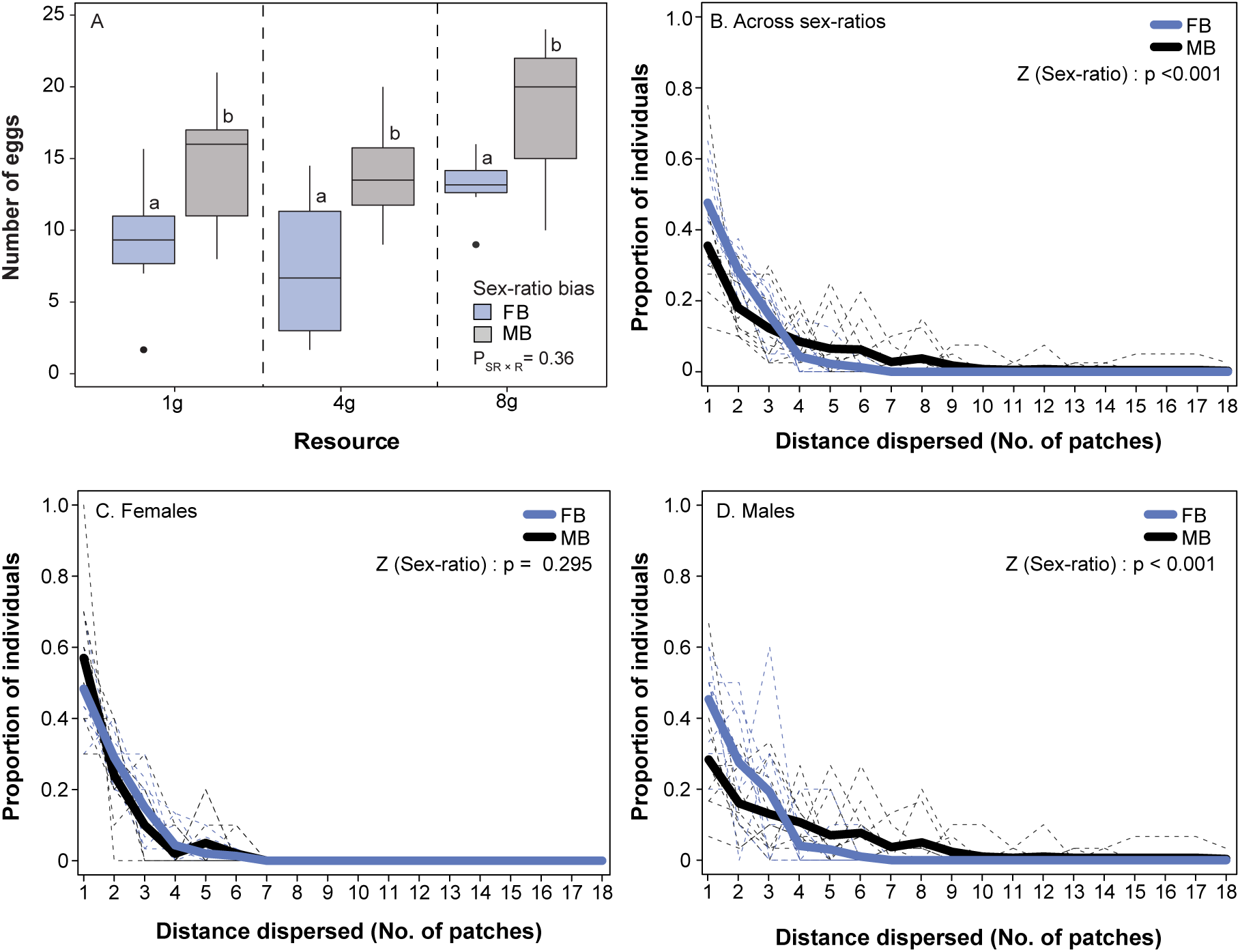
(A) Fecundity as a function of sex-ratio bias (i.e. Male-biased: MB vs Female-biased: FB) and natal resource (wheat flour) availability (Data analysed using an ANCOVA); (B-D) Effects of sex-ratio bias on dispersal. Dispersal kernels for individuals from different treatment combinations across sexes (males vs females) and sex-ratio groups (MB vs FB): Comparison across (B) sex-ratio groups; (C) females and (B) males separately across sex-ratio groups (n=10 replicates/sex-ratio bias). In each panel, dotted lines represent the proportion of beetles found in different patches of individual linear arrays across sexes and sex-ratio groups; whereas bold lines represent their means across all linear arrays. In panel A, P-value represents the result of the interaction between sex ratio bias (SR) and resource availability (R). In panel A, significantly different groups are connected with different alphabets (based on Tukey’s HSD), and alphabet assignments are meaningful only within each resource level and are not comparable across resource levels. In panels B-D, P-values represent results from the ordinal logistic model best fitted with the logit link function.

### Females did not disperse to avoid interference competition

We then tested whether increased dispersal can be an alternative mechanism for females to avoid interference competition. We found significant main effects of both sex and group sex-ratio (**Fig.** 2B-D, S3A-B; **Table** S2A). On average, while MB beetles (as a group) were 44.52% more likely to disperse than FB beetles, individual males were 54.94% more likely to disperse than females (**Fig.** S3A). MB groups or males also dispersed over a greater distance than FB beetles and females respectively (significant effects on mean, standard deviation, maximum), while the shape of the dispersal kernel did not change across sexes and sex-ratios (no significant impact on skew or kurtosis, **Fig.** S3C-E; **Table** S2B). However, a significant interaction between sex and sex-ratio revealed that the observed effects were primarily driven by highly dispersive MB males, whereas no significant difference was found in dispersal between MB and FB females (**Fig.** 2C vs D, S3A; **Table** S2). This contradicted our expectations that FB females would disperse more than their MB counterparts to avoid competition.

We also confirmed that although quinones mediated female competition at high female-density (Khan *et al.* 2018), they might not serve as a chemical cue for female dispersal─ females did not disperse more when they were exposed to a high concentration of MBQ in their natal habitat (also see SI results; **Fig.** S4, **Table** S3).

### Carcass scavenging relaxed the female interference competition

We next tested whether scavenging of carcasses could relax the competition. Overall, FB groups showed higher scavenging activity than MB groups (**Fig.** 3A, **Table** S1B), but the effect was likely driven by collective effects of high female density within FB groups, with females typically showing higher per capita scavenging than males (**Fig.** 3B, **Table** S1C). Surprisingly, when assayed individually after removing the sex-ratio bias (see methods), MB females scavenged more than FB females, possibly because MB females required more resource supplement to maintain their higher reproductive output (**Fig.** 3B, **Table** S1C). However, despite these contrasting variations across sexes and sex-ratio groups, scavenging led to ~2-fold rise in fecundity of FB females (**Fig.** 3C, **Table** S1D). We also found that scavenging significantly reduced the zone of inhibition produced by individual cold-shocked FB females, indicating a reduced quinone production inside stink glands (**Fig.** 3D, **Table** S1E). Interestingly, scavenging did not impact fecundity or quinone content of MB females as they were perhaps already reproducing at an optimal rate (**Fig.** 3C-D, **Tables** S1D-E). Overall, these results suggested a drastic reduction of interference competition between FB females after scavenging, with increased fecundity and reduced chemical warfare.

**Fig 3.**
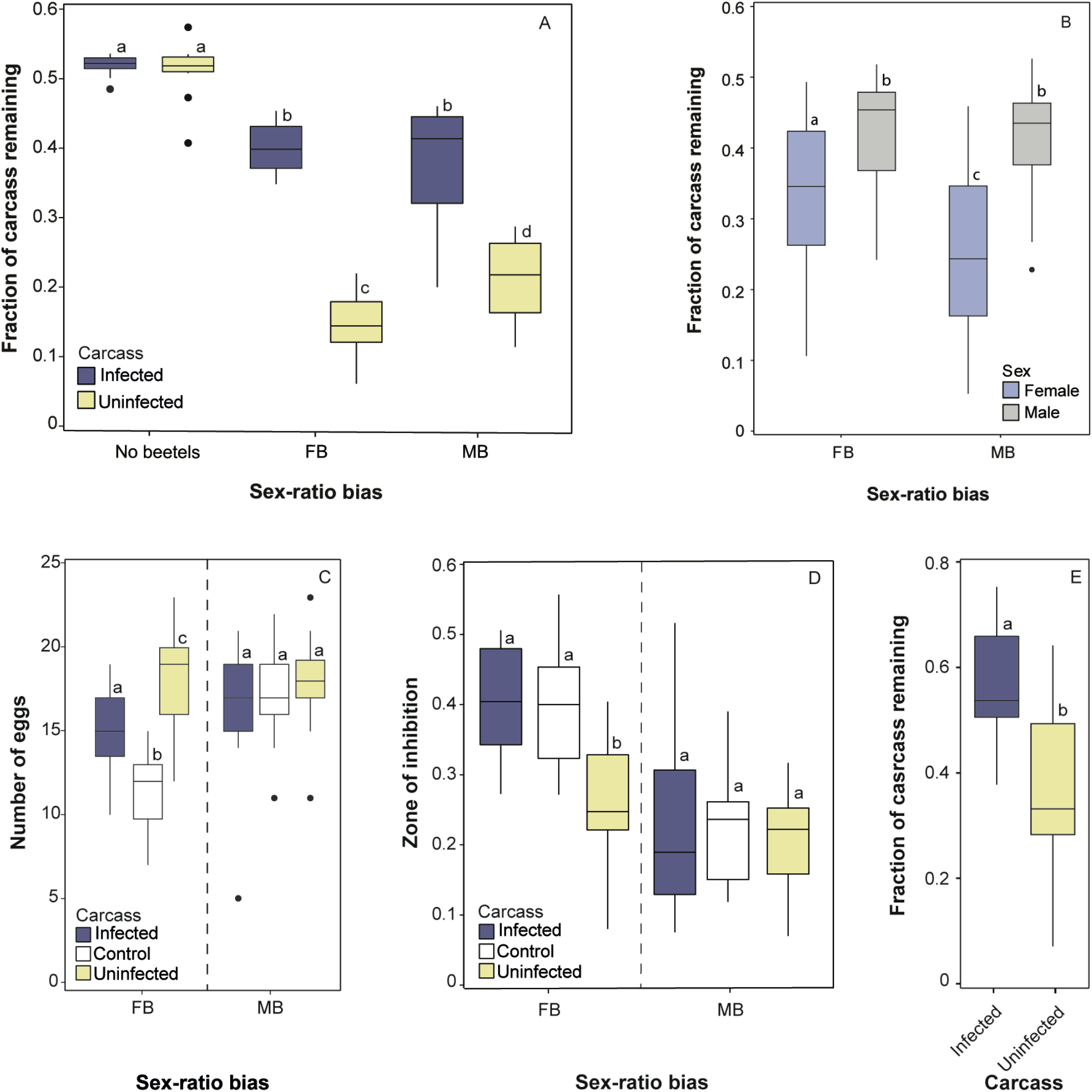
Effects of scavenging. (A) Scavenging activity as a function of carcass type (i.e. normal or infected carcass or no beetles) and sex-ratio bias (Male-biased: MB vs Female-biased: FB) (Data analysed using a generalised linear model best fitted to quasibinomial distribution with logit link function); (B) Effect of sex-ratio bias on the scavenging activity of individual beetles (Data analysed using a generalised linear model best fitted to quasibinomial distribution with logit link function) (n= 25 beetles/sex/sex-ratio); (C) Fecundity as a function of sex-ratio bias and carcass type (i.e. normal or infected carcass or no carcass) (Data analysed using a two-way ANOVA); (D) Zone of inhibition produced by individual females as a function of sex-ratio bias and carcass type (Data analysed using a two-way ANOVA); Y-axis represents normalised ZIs produced by each beetle (described in **Fig.** 1); (E) Two-way choice of scavenging as a function of carcass type (n= 13 replicates/carcass type) (Data analysed using a paired t-test). Replicate sizes for experimental results described in panels A, C-D have been reported in Fig 1; In panels A-E, significantly different groups are connected with different alphabets (based on Tukey’s HSD). In panels C-D, alphabet assignments are meaningful only within each sex-ratio group (partitioned by dashed vertical lines) and are not comparable across sex-ratio groups.

### Scavenging was beneficial, regardless of the carcass quality

Animal death due to infection and disease can produce considerable variation in the quality of available carcasses (Devault *et al.* 2003; Sage *et al.* 2019). Infected carcasses may have lower nutritional value due to microbial overgrowth (Devault *et al.* 2003) and increased infection risk (Sage *et al.* 2019), reducing the net fitness effects of scavenging. To test this possibility, we directly manipulated the quality of carcasses by infecting larvae with a lethal dose of Bt and then quantified the scavenging and fitness effects. When given a simultaneous choice between infected vs normal carcasses, individual beetles showed lower scavenging of infected carcasses (**Fig.** 3E, **Table** S1F), indicating an intrinsic avoidance. However, despite the bias, we noticed considerable scavenging of infected carcasses in the absence of choice (**Fig.** 3A, **Table** S1B) and competing FB females were able to increase their fecundity more than their control counterparts (**Fig.** 3C, **Table** S1D). Conversely, we also found that the scavenging activity of infected carcasses and its fitness benefits were lower than that of uninfected carcasses in FB females (**Fig.** 3C, **Table** S1D), highlighting the apparent costs of scavenging infected carcasses. Interestingly, we did not find any impact of scavenging infected carcasses on stink gland production of FB females (Fig. 3D, **Table** S1E). Hence, fitness effects of scavenging infected carcasses might strongly depend on specific ecological contexts─ e.g. although they might appear less-preferred and costlier than scavenging standard uninfected carcasses, they can still remain highly beneficial under competitive conditions.

### Protein supplement reduced female interference competition

Since female interference competition was reduced by carcass scavenging but not by the *ad libitum* access to the natal resource, we suspected more specific roles of carcass-derived proteins (Huntington & Higley 2010). To verify this hypothesis, we compared the fecundity of females from different sex-ratio groups that had the opportunity to scavenge *T. castaneum* larvae versus those who fed on (natal resource) wheat flour supplemented with dry yeast, a known protein source that boosts fecundity in insects (Matzkin *et al.* 2011). As expected, FB females again scavenged more than MB females (**Fig** 4A, **Table** S4A). Both carcass scavenging and yeast supplementation produced an equivalent increase in fecundity of FB females than their control groups (**Fig** 4B, **Table** S4B), highlighting the likely role of proteins in the observed fitness recovery. We also found similar fitness benefits with scavenging of larval carcasses derived from another beetle *T. molitor*, suggesting a more generalised role of acquiring animal proteins through carcass scavenging during interference competition, rather than cannibalism of conspecific larvae (**Fig** 4A-B, **Table** S4A-B). None of these treatments had any impact on the fecundity of MB females.

**Fig 4.**
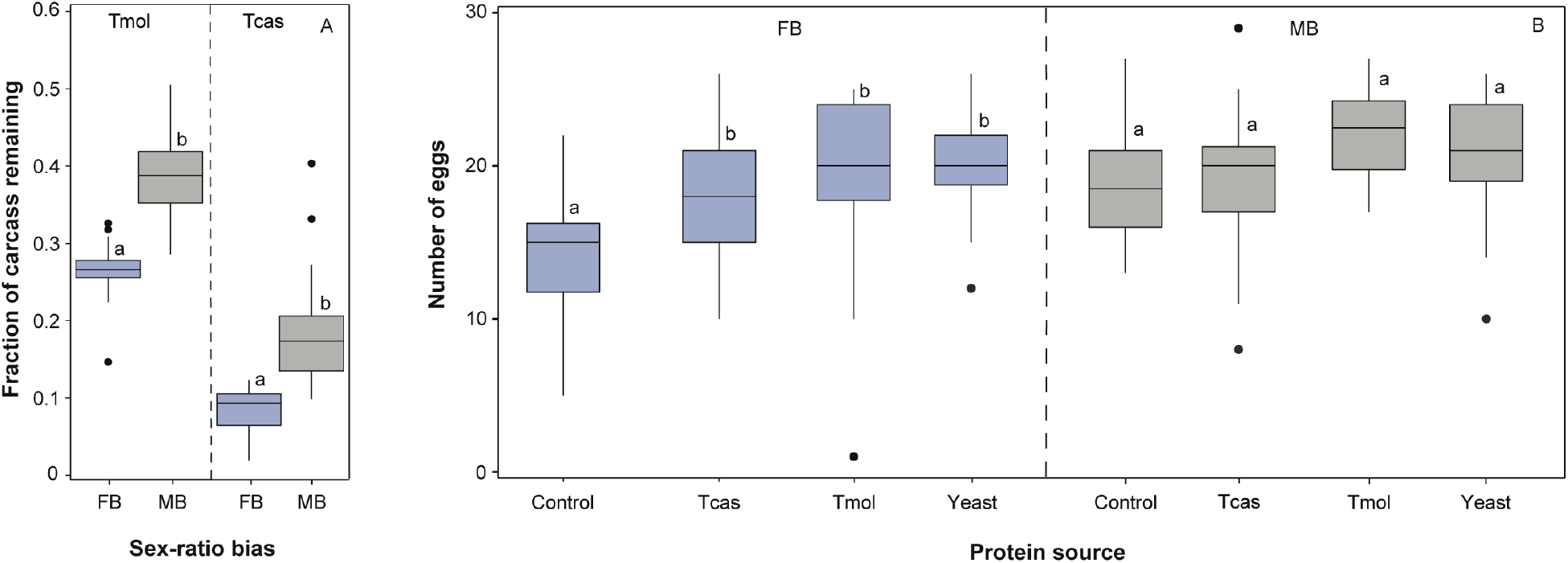
Comparing the effects of scavenging and dry yeast as a protein supplement during female interference competition. (A) Scavenging activity as a function of the carcass from different animals (*T. castaneum*; Tcas or *T. molitor;* Tmol) and sex-ratio (Male-biased: MB vs Female-biased: FB) (n= 19-20 replicates/protein source/sex-ratio). Scavenging data was analysed using a generalised linear model, as described in **Fig.** 3. (B) Fecundity as a function of protein source (i.e. carcasses from different animals, yeast or control) and sex-ratio (n= 19-20 replicates/protein source/sex-ratio). Fecundity data was analysed using a generalised linear model, fitted to Poisson distribution. Significantly different groups are connected with different alphabets (based on Tukey’s HSD). Alphabet assignments are meaningful only within each type of animal carcass (or sex-ratio) (partitioned by dashed vertical lines). They are not comparable across types of animal carcasses (or sex-ratio groups).

### Stink gland contents triggered scavenging to increase fecundity

Finally, we tested whether the accumulation of stink gland products can serve as chemical cues to regulate scavenging and fitness effects. Indeed, we found that exposure to abdominal stink gland extracts increased scavenging in previously mated test females (**Fig** 5A, **Table** S5A). While stink gland exposure generally reduced female fecundity (also see Khan *et al.* 2018), access to the carcass and increased scavenging triggered by stink gland extracts helped females to overcome the negative fitness effects of stink gland exposure and maintain their fecundity as high as the respective control group (**Fig** 5B, **Table** S5B).

Subsequently, we identified two major stink gland components such as MBQ and PD which independently mirrored the effects of whole stink gland extracts (**Fig** 5C-D, **Table** S5A-B), suggesting their functional role in increased scavenging and the observed fitness effects. Taken together, these data suggested an exciting possibility of a feedback loop where the same stink gland chemicals (e.g. MBQ) that reduced female fitness during interference competition (Khan et al. 2018) can also serve as a potential catalyst for increased scavenging to compensate the fitness loss.

**Figure 5.**
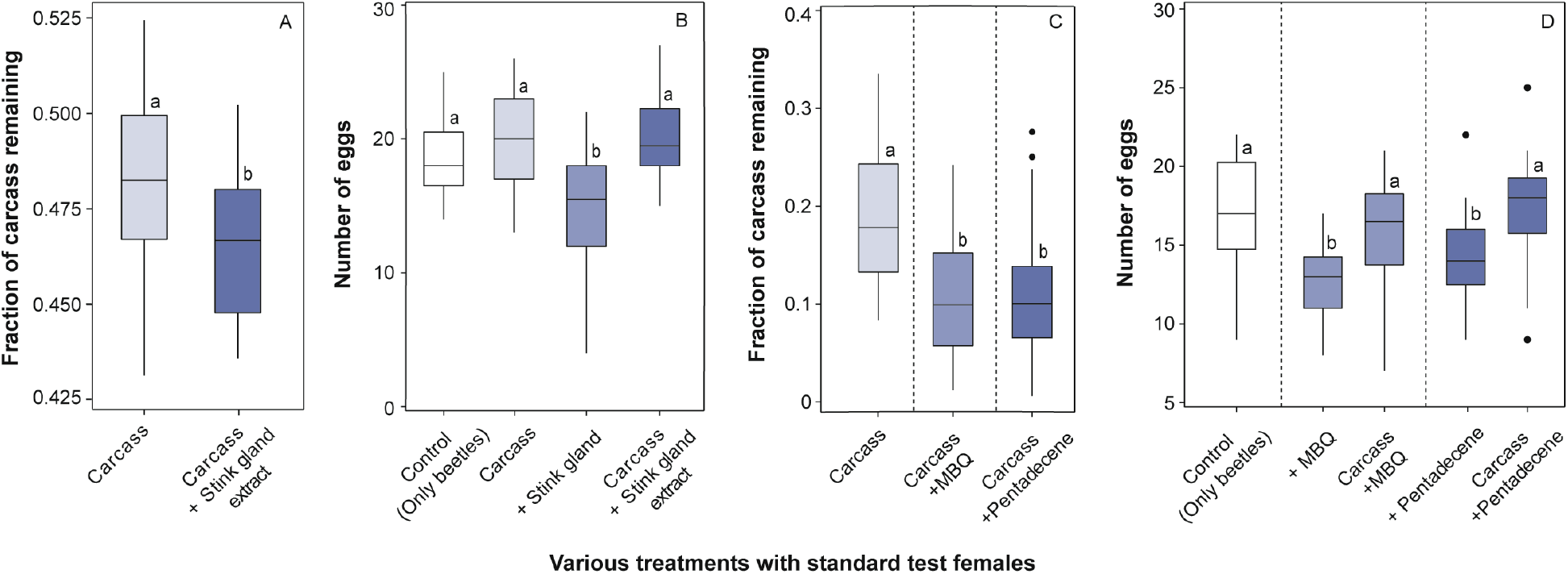
Effect of stink gland contents on scavenging activity and fitness effects. (A) Scavenging activity and (B) fecundity of standard test females, after exposure to whole abdominal stink glands extracted in hexane (n= 18-19 replicates/assay/treatments). (C) Scavenging activity and (D) fecundity of standard test females, after exposure to individual stink gland components such as methyl-p-benzoquinone (MBQ) and 1-pentadecene (PD) dissolved in hexane (n= 20 replicates/assay/treatments). Scavenging data was analysed using a generalised linear model, as described in **Fig**. 3. Fecundity data was analysed using a one-way ANOVA. Only solvent (Hexane) did not have any impact on fecundity (negative control) (ANOVA: P= 0.9). Significantly different groups are connected with different alphabets (based on Tukey’s HSD). In panels C-D, alphabets linking MBQ or PD treatment (partitioned by the dotted line) can be compared separately with the control treatment. They are not comparable between MBQ and PD treatments.

## DISCUSSION

Despite the long-standing interests in the roles of facultative scavenging in interspecific competition and ecological food web (Wilson & Wolkovich 2011; Moleón *et al.* 2014), we have relatively little understanding of how it might affect intra-specific resource competition between females. This is not surprising as most studies interpret intrasexual competition in the light of male-male competition for mating, thereby limiting our scope of analysing female-female resource competition. However, the surge of evidence, in recent years, for female intra-sexual competition, particularly in mammals, for food and space by dominance relationships, territoriality and inter-group aggression suggests that it is perhaps more widespread than what was assumed previously (reviewed in Stockley & Bro-Jørgensen 2011). A new study found female competitions even in insects (Khan *et al.* 2018)─ e.g. *Tribolium* females inhibited each other’s reproduction by releasing benzoquinones in response to the increased female density of female-biased (FB) groups, suggesting chemical interference competition. In the present work, we clearly showed that the fitness effects of such interference competition in FB beetles could be entirely reversed by allowing them to scavenge on larval carcasses. Scavenging not only increased fecundity of FB females but also reduced the production of stink gland contents, suggesting overall relaxation of interference competition. Although previous studies reported the importance of carcass access during the intraspecific female competition in obligate scavengers such as burying beetles (Trumbo 1990), our work is perhaps first to experimentally demonstrate that carcass availability could relax chemical interference competition in facultative scavengers. Since both sex-ratio and female density vary widely across species and populations, we expect that carcass-derived fitness benefits might have strong implications for natural populations as well during female interference competition.

Interestingly, although dispersal is a key ecological process to avoid the competition of high-density groups (Vahsen *et al.* 2018; Endriss *et al.* 2019), it did not play any specific role in reducing the interference competition between females. Despite the apparent increase in toxic quinone accumulation at a high female density of FB groups (Khan et al. 2018), FB females did not disperse more than MB females to escape the chemical competition. Quinone’s lacking role was further supported by another experiment where groups of mated females did not increase dispersal even after they were directly exposed to a high concentration of MBQ continuously for 36 hours. These results were puzzling because beetles typically avoid flour containing a high level of quinones (Loconti & Roth 1953). We thus speculate that the overall adverse fitness effects of chemical secretions in beetles might still not be sufficient to overcome various unanticipated costs associated with female dispersal (e.g. energetic cost of movement, risk of predations or failure to find a favourable patch to colonise; see Li & Kokko 2019). Alternatively, beetles were already tolerant to fluctuations in quinone levels in their environment, evolving a quinone-mediated density-dependent fecundity regulation to maintain the population growth (Sonleitner & Gutherie 1991). Hence, although quinones can play a critical role in signalling competition and limiting reproduction to regulate population density (Sonleitner & GuthJanis 1991), they might be irrelevant for dispersal to explore new patches. In contrast to females, male dispersal increased significantly at a high male density. This is consistent with existing theories predicting that MB sex-ratios might favour male-dispersal as an adaptive response to reduce intra-sexual competition to find mates and avoid inbreeding (Nelson & Greeff 2011). These results thus also underscore the divergent pattern of dispersal behaviour across sexes during intrasexual competition.

However, one of the most critical findings of this study is that female reproductive competition was regulated by specific nutrients derived from carcasses, but not the total resource available to each female. In contrast to fitness effects of scavenging, increased access to wheat flour (as high as 8-fold), which is also the natural diet for flour beetles, had no impact on fecundity. Since carcasses are highly nutritious (Hansky 1987) and serve as a critical component of the biogeochemical cycling of free amino acids and peptide pool (Macdonald *et al.* 2014), one possibility was that protein intake via carcass feeding was responsible for reducing the level of female interference competition in our experiments. Indeed, we found that active dry yeast, a protein supplement that usually boosts sexual activity and fecundity in insects (Perez-Staples et al. 2007; Matzkin et al. 2011), also mirrored the same fitness effects as that of carcass feeding, increasing the fecundity of FB females. Previous studies in carcass-feeding blowflies supported the hypothesis as well where decomposed flesh served as a vitellogenic protein source to increase reproduction (Huntington and Higley 2010). Together, these results suggested novel aspects of female competition where proteins derived from carcasses might play a more important role than the availability of natal resources (Stockley & Bro-Jørgensen 2011). We note that the observed fitness benefits from scavenging of *Tribolium* larvae can also be interpreted as specific effects of cannibalism of dead conspecifics. Although cannibalism is indeed a highly relevant solution to acquire essential nutrients during the competition (Via 1999; Richardson *et al.* 2010), it is unlikely to explain our results completely because we found similar fitness benefits from scavenging of *T. molitor* carcasses as well, suggesting a more generalised role of scavenging.

Another striking outcome of this study is that scavenging was modulated by specific stink gland components, revealing its novel chemical regulations and fitness effects. Exposure to total gland extracts, as well as individual components such as MBQ (Khan et al. 2018) and PD, reduced egg production in standard focal females, but they also increased scavenging upon the availability of larval carcasses, with a correlated increase in fecundity. Based on these results, we propose that density-dependent accumulation of stink gland secretion is not just a reliable indicator of female interference competition (Khan *et al.* 2018), but might also serve as a cue to adopt effective strategies such as scavenging to resolve the resource conflicts. We thus propose a novel feedback loop between stink gland components, scavenging and fitness that might regulate the outcome of female competitive interactions. Feedback loops may have strong implications for natural beetle populations as well because skewed sex-ratios, high density and availability of carcasses due to high juvenile mortality might frequently co-occur in the wild.

We also identified carcass quality, such as infection status, as an essential ecological parameter that might affect the propensity of scavenging and fitness effects during interference interaction. When given a choice, beetles strongly biased against feeding on larval carcasses that died of Bt infection. We also noted a significant reduction in group scavenging of infected carcasses, with a concomitant decline in fecundity compared to control females fed on uninfected carcass. Yet females fed with infected carcasses showed higher fertility than control FB groups (with no available carcasses), highlighting the potential benefits of scavenging during the competition, regardless of the carcass quality. We speculate that the risk of transmitting bacterial infection can also be high with infected cadavers (Unpublished data, Shreya Verma; Milutinović *et al.* 2015). However, any such costs of infection transmission were perhaps outweighed by the immediate fitness benefits accrued via protein acquisition during competitive interactions. Also, an interesting observation was that scavenging of infected carcasses did not reduce quinone production in FB females (compared with scavenging of normal carcass). One possibility is that due to the potential risk of infection, females did not reduce their stink gland components which can act as externally secreted antimicrobials to clean the surrounding environment (Joop *et al.* 2014; Khan *et al.* 2015), thereby preventing Bt transmission. Since a large proportion of available carcasses in the wild originate from diseased animals (Brand 2013), thereby increasing the infection risk and fitness costs (Pearman *et al.* 2004; Rebolledo *et al.* 2015), we think these results are highly relevant to natural conditions as well.

In closing, we want to highlight that among several plausible ecological contexts that we tested, scavenging of proteinaceous carcass emerged as the most relevant strategy to overcome the chemical interference competition in female flour beetles. We hope that the suggested role of a reciprocal feedback loop between scavenging, stink gland chemicals and fitness will motivate future work to trace detailed chemical ecology involved in the detection and processing of such information during competitive interactions. Our results, revealing the life-history contexts (e.g. female competition) that render even scavenging of infectious carcasses highly beneficial, might instigate new research questions to analyse the ecological conditions where disease and infection can spread via scavenging in the wild. Finally, as more evidence is accumulating for female interference competition from various species (Stockley & Bro-Jørgensen 2011), our results might serve as a rare systematic template to advance the future understanding of vital ecological determinants that influence female competition.

## ACKNOWLEDGEMENTS

We are grateful to Debapriyo Chakraborty, Deepa Agashe and Shivani Krishna for their feedback on the manuscript. We are particularly thankful to Swastika Issar, Arun Prakash and Deepa Agashe for their help and inputs in the experiment measuring the impact of natal resource on beetle fecundity. We also thank Shreya Verma and Devshuvam Banerji for their support during experiments and Saubhik Sarkar for providing the beetle image.

## AUTHOR CONTRIBUTIONS

IK and BB conceived and designed experiments; BB, MR, SS and IK carried out experiments; BB, DNB and IK analysed data; IK acquired funding; IK wrote the manuscript with inputs from BB, SS, MR and DNB. All authors gave final approval for publication.

## REFERENCE

Abdel-Razek, A.S., Salama, H.S., White, N.D.G. & Morris, O.N. (1999). Effect of *B*acillus *thuringiensis* on feeding and energy use by *Plodia interpunctella* (Lepidoptera: Pyralidae) and *Tribolium castaneum* (Coleoptera: Tenebrionidae). Can. Entomol., 131, 433–440.

Abernathy, E.F., Turner, K.L., Beasley, J.C. & Rhodes, O.E. (2017). Scavenging along an ecological interface: utilization of amphibian and reptile carcasses around isolated wetlands. Ecosphere, 8:e01989.

Alabi, T., Michaud, J.P., Arnaud, L. & Haubruge, E. (2008). A comparative study of cannibalism and predation in seven species of flour beetle. Ecol. Entomol., 33, 716–726.

Bell, M.B.V., Cant, M.A., Borgeaud, C., Thavarajah, N., Samson, J. & Clutton-Brock, T.H. (2014). Suppressing subordinate reproduction provides benefits to dominants in cooperative societies of meerkats. Nat. Commun., 5, 4499.

Borchering, R.K., Bellan, S.E., Flynn, J.M., Pulliam, J.R.C. & McKinley, S.A. (2017). Resource-driven encounters among consumers and implications for the spread of infectious disease. J. R. Soc. Interface, 14, 20170555.

Brand, C.J. (2013). Wildlife mortality investigation and disease research: contributions of the USGS National Wildlife Health Center to endangered species management and recovery. Ecohealth, 10, 446–454.

Clobert, J., Le Galliard, J.F., Cote, J., Meylan, S. & Massot, M. (2009). Informed dispersal, heterogeneity in animal dispersal syndromes and the dynamics of spatially structured populations. Ecol. Lett., 12, 197–209.

Clutton-Brock, T.H. & Lukas, D. (2012). The evolution of social philopatry and dispersal in female mammals. Mol. Ecol., 21, 472–492.

Corrales-Carvajal, V.M., Faisal, A.A. & Ribeiro, C. (2016). Internal states drive nutrient homeostasis by modulating. eLife, 5:19920, 1–28.

Dawson, P.S. (1977). Life history strategy and evolutionary history of *Tribolium* flour beetles. Evolution., 31, 226–229.

Devault, T.L., Rhodes, O.E., Shivik, J.A. (2003). Scavenging by vertebrates: behavioral, ecological, and evolutionary perspectives on an important energy transfer pathway in terrestrial ecosystems. Oikos, 102, 225–234.

Endriss, S.B., Vahsen, M.L., Bitume, E. V., Grey Monroe, J., Turner, K. G., Norton, A.P., et al. (2019). The importance of growing up: juvenile environment influences dispersal of individuals and their neighbours. Ecol. Lett., 22, 45–55.

Halliday, W.D., Slevan-Tremblay, I. & Blouin-Demers, G. (2019). Do female red flourbeetles assess both current and future competition during oviposition? J. Insect Behav., 32, 181–187.

Slansky, F. & Rodriguez, J. G. (1987). Nutritional ecology of insects, mites, spiders, and related invertebrate. Wiley NY.

Harvey, J.A., Poelman, E.H. & Tanaka, T. (2013). Intrinsic inter- and intraspecific competition in parasitoid wasps. Annu. Rev. Entomol., 58, 333–351.

Ho, F.K.. & Dawson, P.S. (1966). Egg cannibalism by *Tribolium* larvae. Ecology, 47, 318–322.

Huntington, T.E. & Higley, L.G. (2010). Decomposed flesh as a vitellogenic protein source for the forensically important *Lucilia sericata* (Diptera: Calliphoridae). J. Med. Entomol., 47, 482–486.

Isles, A.R., Baum, M.J., Ma, D., Szeto, A., Keverne, E.B. & Allen, N.D. (2002). A possible role for imprinted genes in inbreeding avoidance and dispersal from the natal area in mice. Proc. R. Soc. B., 269, 665–670.

Joop, G., Roth, O., Schmid-Hempel, P. & Kurtz, J. (2014). Experimental evolution of external immune defences in the red flour beetle, J. Evol. Biol., 27, 1562–1571

Khan, I., Prakash, A. & Agashe, D. (2015). Immunosenescence and the ability to survive bacterial infection in the red flour beetle *Tribolium castaneum*. J. Anim. Ecol., 85, 291–301.

Khan, I., Prakash, A. & Agashe, D. (2017). Experimental evolution of insect immune memory versus pathogen resistance. Proc. R. Soc. B., 284, 20171583.

Khan, I., Prakash, A., Issar, S., Umarani, M., Sasidharan, R., Masagalli, J.N., et al. (2018). Female density-dependent chemical warfare underlies fitness effects of group sex ratio in flour beetles. Am. Nat., 191, 306–317.

Latham, N. & Mason, G. (2004). From house mouse to mouse house: the behavioural biology of free-living *Mus musculus* and its implications in the laboratory. Appl. Anim. Behav. Sci., 86, 261–289.

Li, J., Lehmann, S., Weißbecker, B., Ojeda Naharros, I., Schütz, S., Joop, G., et al. (2013). Odoriferous defensive stink gland transcriptome to identify novel genes necessary for quinone synthesis in the red flour beetle, *Tribolium castaneum*. PLoS Genet., 9, e1003596.

Li, X.Y. & Kokko, H. (2019). Sex-biased dispersal: a review of the theory. Biol. Rev., 94, 721–736.

Loconti, J.D. & Roth, L.M. (1953). Composition of the odorous secretion of *Tribolium castaneum*. Ann. Entomol. Soc. Am., 46, 281–289.

Macdonald, B.C.T., Farrell, M., Tuomi, S., Barton, P.S., Cunningham, S.A. & Manning, A.D. (2014). Carrion decomposition causes large and lasting effects on soil amino acid and peptide flux. Soil Biol. Biochem., 69, 132–140.

Matthysen, E. (2005). Density-dependent dispersal in birds and mammals. Ecography, 28, 403–416.

Matzkin, L.M., Johnson, S., Paight, C., Bozinovic, G. & Markow, T.A. (2011). Dietary protein and sugar differentially affect development and metabolic pools in ecologically diverse *Drosophila*. J. Nutr., 141, 1127–1133.

Mckillup, S.C. & Mckillup, R. V. (1996). The feeding behaviour of *Thalamita crenata* (Portunidae, Decapoda), a cannibalistic marine scavenger. Mar. Freshw. Behav. Physiol., 28, 255–267.

Michalczyk, Ł., Millard, A.L., Martin, O.Y., Lumley, A.J., Emerson, B.C. & Gage, M.J.G. (2011). Experimental evolution exposes female and male responses to sexual selection and conflict in *Tribolium castaneum*. Evolution, 65, 713–24.

Milutinović, B., Höfling, C., Futo, M., Scharsack, J.P. & Kurtz, J. (2015). Infection of *Tribolium castaneum* with *Bacillus thuringiensis*: quantification of bacterial replication within cadavers, transmission via cannibalism, and inhibition of spore germination. Appl. Environ. Microbiol., 81, 8135–8144.

Moleón, M., Sánchez-Zapata, J.A., Selva, N., Donázar, J.A. & Owen-Smith, N. (2014). Inter-specific interactions linking predation and scavenging in terrestrial vertebrate assemblages. Biol. Rev., 89, 1042–1054.

Murray, C.M., Eberly, L.E. & Pusey, A.E. (2006). Foraging strategies as a function of season and rank among wild female chimpanzees (*Pan troglodytes*). Behav. Ecol., 17, 1020–1028.

Nelson, R.M. & Greeff, J.M. (2011). Sex ratio dependent dispersal when sex ratios vary between patches. J. Theor. Biol., 290, 81–87.

Ogden, J.C. (1969). Effect of components of conditioned medium on behavior in *Tribolium confusum*. Physiol. Zool., 42, 266–274.

Pearman, P.B., Garner, T.W.J., Straub, M. & Greber, U.F. (2004). Response of the Italian agile frog (*Rana latastei*) to a ranavirus, frog virus 3: a model for viral emergence in naïve populations. J. Wildl. Dis., 40, 660–669.

Perez-Staples, D., Prabhu, V. & Taylor, P.W. (2007). Post-teneral protein feeding enhances sexual performance of Queensland fruit flies. Physiol. Entomol., 32, 225–232.

Piper, M.D.W., Blanc, E., Leitão-Gonçalves, R., Yang, M., He, X., Linford, N.J., et al. (2014). A holidic medium for *Drosophila melanogaster*. Nat. Methods, 11, 100–105.

Porter, J.H. & Dooley Jr., J.L. (1993). Animal Dispersal Patterns: A reassessment of simple mathematical models. Ecology, 74, 2436–2443.

Pusceddu, M., Mura, A., Floris, I. & Satta, A. (2018). Feeding strategies and intraspecific competition in German yellowjacket (*Vespula germanica*). PLoS One, 13, e0206301.

Rebolledo, D., Lasa, R., Guevara, R., Murillo, R. & Williams, T. (2015). Baculovirus-induced climbing behavior favors intraspecific necrophagy and efficient disease transmission in *Spodoptera exigua*. PLoS One, 10, e0136742.

Richardson, M.L., Mitchell, R.F., Reagel, P.F. & Hanks, L.M. (2010). Causes and consequences of cannibalism in noncarnivorous insects. Annu. Rev. Entomol., 55, 39–53.

Sage, M.J. Le Towey, B.D. & Brunner, J.L. (2019). Do scavengers prevent or promote disease transmission? The effect of invertebrate scavenging on *Ranavirus* transmission, Funct Ecol., 33,1342–1350.

Sokoloff, A. (1977). The biology of Tribolium. Oxford, UK Press. Clarendon.

Sonleitner, F.J. & Gutherie, J. (1991). Factors affecting oviposition rate in the flour beetle *Tribolium castaneum* and the origin of the population regulation mechanism. Res. Popul. Ecol. (Kyoto)., 33, 1–11.

Stockley, P. & Bro-Jørgensen, J. (2011). Female competition and its evolutionary consequences in mammals. Biol. Rev., 86, 341–366.

Tobias, J.A., Montgomerie, R. & Lyon, B.E. (2012). The evolution of female ornaments and weaponry: social selection, sexual selection and ecological competition, Philos. Trans. R. Soc. B., 367, 2274–2293.

Trumbo, S.T. (1990). Interference competition among burying beetles (Silphidae, *Nicrophorus)*. Ecol. Entomol., 15, 347–355.

Turner, K.L., Abermethy, E.F., Conner, L.M., Rhodes, O.E. & Beasley, J.C. (2017). Abiotic and biotic factors modulate carrion fate and vertebrate scavenging communities. Ecology, 98, 2413–2424.

Unruh, L.M., Xu, R. & Kramer, K.J. (1998). Benzoquinone levels as a function of age and gender of the red flour beetle, *Tribolium castaneum*, Insect Biochem. Mol. Biol., 28, 969–977.

Vahsen, M.L., Shea, K., Hovis, C.L., Teller, B.J. & Hufbauer, R.A. (2018). Prior adaptation, diversity, and introduction frequency mediate the positive relationship between propagule pressure and the initial success of founding populations. Biol. Invasions, 20, 2451–2459.

Via, S. (1999). Cannibalism facilitates the use of a novel environment in the flour beetle, *Tribolium castaneum*, Heredity, 82, 276–275.

Waser, P.M. (1985). Does competition drive dispersal ? Ecology, 66, 1170–1175.

Wheeler, D. (1996). The role of nourishment in oogenesis. Annu. Rev. Entomol. Vol. 41, 407–431.

Wilson, E.E. & Wolkovich, E.M. (2011). Scavenging: how carnivores and carrion structure communities. Trends Ecol. Evol., 26, 129–135.

