## Supplementary Information for "Carcass Scavenging Relaxes Chemical-Driven Female Interference Competition"

^1^Ashoka University

Bangalore 560065, India

**1. SUPPLEMENTARY METHODS**

**Generating experimental individuals**

We used an outbred laboratory population of *Tribolium castaneum* for all the experiments. We maintained the population on whole-wheat flour at 32°C, with a 45-day discrete generation cycle. To generate the experimental individuals, we allowed ~1000 adult beetles to oviposit on 400gm of wheat flour for 24 hours. We collected the virgin offspring as pupae after three weeks and sexed them. We isolated these pupae individually into the wells of 96-well microplates for 14 days post-pupation. Since the pupal stage usually lasts for 3–4 days in our beetle population, we thus obtained ∼10-day-old (post-eclosion), sexually mature virgin adults for all the experiments.

**Quantification of dispersal behaviour under biased sex-ratios**

We constituted MB and FB groups with same level of sex-ratio bias as described earlier but with an increased group size of 40 beetles (FB: 10 males + 30 females; MB: 30 males + 10 females) (n=10 replicates/ sex-ratio bias group) to obtain a meaningful comparison between dispersal tendencies across sex-ratio groups. Also, our previous experiments confirmed that the interference competition within FB groups generally reduced female fitness, regardless of the group size of beetles (Khan *et al.* 2018). To perform the dispersal assays, we closely followed the protocol described by (Endriss *et al.* 2019), manufacturing linear arrays of twenty 4 × 4 × 6 cm^3^ plastic boxes (or patches) that were connected by 2mm holes. Since none of our pilot studies found beetles from stock populations dispersing more than 18 patches, we thus assumed that beetle dispersal was not limited by the length of the array. We first added beetles to the linear array and allowed them to acclimatise in the first patch for 48 hours by blocking the holes connecting the 1^st^ and 2^nd^ patch, with plastic plugs. After the acclimatisation period, we removed the plugs to allow beetle groups to disperse among patches. We stopped the dispersal after 30 hours and immediately counted all experimental males and females within each patch across linear arrays. This protocol enabled us to accurately estimate how many patches they dispersed from their patch of origin (i.e. the first patch within the array). Since the total number the males and females differed between sex-ratio groups, we transformed the census data as the fraction of total number of males or females actually present in their respective sex-ratio groups (e.g. if there are 5 FB females in any given patch of linear arrays, we transformed the value as 5/30= 0.167), as the unit of analysis for dispersal distance.

**Quantification of dispersal behaviour as a function of quinones**

To directly quantify dispersal tendencies as a function of chemical cues provided by quinones, we used almost a similar protocol as described above. We used 10-day-old virgin beetles to constitute 10 mating pairs and added them to the first patch of the linear arrays described earlier (see Fig 1) (n= 9 replicates/treatment). After an initial acclimatisation period of 48 hours, we added filter paper discs soaked in 100µg of commercially available methyl-benzoquinone (MBQ) dissolved in hexane (or only hexane as procedural control) and allowed beetles to disperse for 36 hours. Subsequently, we counted and sexed the individuals in each patch.

**Per capita scavenging as a function of sex-ratio groups**

To tease apart the effects of group scavenging with unequal sex-ratios versus per capita consumption, we quantified the scavenging rate of individual beetles from MB and FB groups (n=25 replicates/sex/sex-ratio bias). To this end, we first constituted biased sex-ratio groups, as described in the main text. After 8 days, we isolated each male and female individually into a 35mm petriplate with 0.5gm flour and then added 4 larval carcasses to each plate to measure scavenging rate as outlined in the main text.

**Zone of inhibition assay, as a proxy for beetle quinone production**

To measure the quinone production, we followed the protocols described earlier (Unruh *et al.* 1998; Prendeville & Stevens 2002; Khan *et al.* 2015), measuring the zone of inhibition (ZI) produced by individual cold-shocked (-80°C for 20 minutes) females embedded in a lawn of bacterial growth on LB agar plates (n = 12-19 females/sex-ratio/carcass type). We first prepared 0.6% LB agar and allowed it to cool down to 45°C degree. Following this, we added freshly grown *Bacillus thuringiensis* culture of 1OD to adjust the final concentration to 0.1OD. After agar plates solidified, we placed individual cold-shocked females to agar plates, with their abdomen embedded in the soft agar. Cold shock induced the complete release of stink gland quinones which diffused into the bacterial lawn on agar plates and produced clear ZI due to strong antimicrobial effects. We added 1µl of antibiotic Streptomycin (100mg/ml) as a positive control to each plate. We incubated the plates at 30°C for 8 hours and then measured the zone of inhibition. We used the mean value of the ZIs measured both horizontally and vertically, as a proxy for quinone content. Further, we normalised ZIs produced by each female, by dividing them with ZIs produced by the antibiotic control in their respective plates, as the unit of analysis. This enabled us to minimise the variations across plates during experimental handling.

**Innate avoidance for scavenging of infected carcasses**

To test whether beetles showed innate avoidance to infected carcasses, we performed a two-way choice experiment─ females were simultaneously provided access to normal vs infected carcass (see Fig S1 for experimental design) and assayed for their scavenging of each carcass type (n=13 replicates/carcass type). We first collected 10-day-old virgin females and mated them with virgin males for 48 hours. We then grouped three 12-day-old females in 35mm Petri-plates with a very thin layer of flour for 8 days, where they had the opportunity to scavenge both infected and uninfected carcasses simultaneously (n=3 larval carcass/infection status/group). To prevent mixing of different carcass types by beetle movement, we immobilised infected and uninfected carcasses separately by impaling them with a 0.1mm insect pin (Fine Science Tools) (See **Fig** S1). To estimate the scavenging activity, we weighed each pin with larvae before and after the experimental window of 8 days as described above.

**Impacts of stink gland components on variation in scavenging rate and fitness effects**

We paired 10-day-old virgins (1 male + 1 female) for 2 days and then isolated the females individually into a 35mm Petri dish with 2gm of flour. On the same day, we also dissected abdominal stink glands from 60 females and prepared larval carcasses as described in the main text. We first homogenised stink glands together in 600μl hexane as described previously in Khan *et al.* 2018. We then soaked filter paper discs of 10 mm diameter with 10μl of supernatant (equivalent to gland extract of 1 female) (or hexane as procedural control). After keeping them under laminar air-flow for 15 minutes to evaporate the solvent, we immediately placed the discs at the centre of Petri plates containing the experimental female. We further divided these experimental females into two subgroups, where females in each plate either received 5 larval carcasses or had no opportunity to scavenge (*n*= 18-19 replicates/treatment/scavenging opportunity). We held these females for 8 days in their respective plates, replacing paper discs soaked with freshly prepared gland extract every 3 days. This was likely to maintain the persistent effects of chemical cues for the entire duration of the experiment. We quantified the scavenging activity and fecundity after as described earlier (n= 18-19 females/treatment/scavenging opportunity)

We followed a similar protocol to test the impacts of individual stink gland components (e.g. MBQ and PD) as well. The only exception was that we placed groups of two mated test females (generated as described above) in each plate to detect changes in the overall scavenging activity more clearly. We exposed these females, with or without the access to the carcass, to a filter paper disc soaked with 15μg MBQ or 5μg PD dissolved in 10μl hexane. We used these concentrations since they were within the physiological range of MBQ and PD content of a single *Tribolium* female (Li *et al.* 2013), mimicking natural interactions. We replaced filter papers with chemicals every three days and quantified scavenging and fecundity after 8 days as described above (n= 20 replicates/treatment/scavenging opportunity).

**Data analysis**

We performed all the analysis in R (version 3.6.3). We tested for the normality of data using the Shapiro-Wilk test, wherever applicable. Model specifications of all the analyses have been described in the supplementary Tables.

Natal resource on the female competition:

We used ANCOVA to analyse the fecundity data as a function of sex-ratio and resource per beetle.

Dispersal as a function of sex-ratio bias and MBQ exposure

1. Dispersal behaviour under biased sex-ratios: We used the transformed data (see SI methods) from each patch across replicate arrays to determine the distribution of dispersed individuals across the linear array (dispersal kernel). We first estimated the differences in dispersal kernels separately across various possible treatment combinations (e.g. MB vs FB groups, Male vs female, MB vs FB females, MB vs FB Males), by fitting the transformed data to an ordinal regression model using *clm* function in the ‘ordinal’ package (Christensen 2019). Subsequently, as described in Endriss *et al.* (2019), we also fitted the transformed data to an ordinal regression model with dispersal distance as the response variable to characterise the overall distribution of beetles from different sexes and sex-ratio groups across linear arrays, using the *clmm* function in the ‘ordinal’ package (Christensen 2019). We used sex-ratio group and sex as fixed effects, with replicate array as a random effect in the linked mixed effect model to account for the orthogonality among individuals within arrays [Model: Distance dispersed ~ Sex-ratio × Sex + (1|replicate array)]. To evaluate the significance of the model terms, we carried out nested likelihood ratio tests. We estimated marginal means of dispersal probability across different treatment combinations using *emmeans* function in the ‘emmeans’ package (Lenth 2020) to evaluate the decumulative probability of dispersal from one patch to the next patch.

We also analysed five parameters of each dispersal kernel such as mean, standard deviation, skew, kurtosis and maximum, using a linear mixed-effects model, with similar fixed and random effect terms as described above. We averaged the model results for each parameter from 1000 simulated runs, by randomly drawing 10 individuals (i.e. the minimum number of individuals/sex/sex-ratio bias in each replicate linear array) per sex from each linear array (Table S4), using *lmer* function in ‘lme4’ package (Bates *et al.* 2015). In other words, we fitted each parameter as response variable from each iteration of simulated, size-independent data to a linear mixed-effects model with sex-ratio group and sex as fixed effects, with replicate array as a random effect (also see Table S4). To generate the simulated dataset, we randomly sampled 10 males and 10 females (total of 20 individuals) from each replicate of MB and FB groups. We then estimated kernel parameters for each replicate and fitted with a mixed-effects model as mentioned earlier for 1000 iterations. Subsequently, we also performed Type II SS ANOVA on the averaged model results from 1000 iterations for each parameter (*car* package, (Fox & Weisberg 2019) (Table S4). Means and confidence intervals for each parameter from the 1000 iterations were estimated using ‘emmeans’ package (Lenth *et al.* 2020).

2. Dispersal behaviour as a function of MBQ exposure: We determined the dispersal kernel as described above. We analysed the data using ordinal regression model using both *clm* and *clmm* function as described above. We used quinone exposure treatments and sex as fixed effects, with replicate array as a random effect. We described the model as [Distance dispersed ~ Quinone exposure treatment × Sex + (1|replicate array)].

Effects of scavenging on the female competition:

1. Scavenging activity and fecundity assay: We analysed the (a) scavenging data (fraction of carcass remaining) using generalised linear model best fitted to quasibinomial distribution (logit link function), with sex-ratio and carcass availability as fixed factors. We also used Hosmer and Lemeshow test to verify the goodness of fit of the model (all P>0.05 ); (b) Fecundity data using a linear mixed-effects model, with sex-ratio and carcass availability as fixed factors and replicate experiment as a random factor. We used Tukey’s HSD to test for pairwise differences after correcting for multiple comparisons, using the R package lsmeans (Lenth 2016).
2. Innate avoidance for scavenging of infected carcasses: We analysed the data using a paired t-test with carcass infection status as a fixed factor.
3. Quinone assay: We analysed the ZI data using a two-way ANOVA with sex-ratio and carcass availability as fixed factors.
4. Analysing the importance of protein supplement during interference competition: We analysed the (a) scavenging data, using a generalised linear model, fitted to quasibinomial distribution, with sex-ratio and protein sources as fixed factors as described above (b) fecundity data, using a generalised linear model, fitted to Poisson distribution, with sex-ratio and food type as fixed factors.

Quantifying the impacts of stink gland extract and individual components on variation in scavenging and fitness effects

We analysed the (a) scavenging data, using a generalised linear model as described earlier (b) fecundity data, using an one-way ANOVA with treatment as a fixed factor. We have analysed the effects of stink gland extract, MBQ and PD separately.

**2. SUPPLEMENTARY RESULTS**

**Exposure to quinones did not affect dispersal behaviour**

Our previous data suggested that exposure to 1µg of MBQ for 12 hours was enough to reduce the fecundity by 25-30% in females (Khan *et al.* 2018). Surprisingly, in this experiment, even 10 µg MBQ per female had no impact on female dispersal tendencies compared to their control counterparts (Fig S4, Table S3). As observed earlier, male dispersed more, regardless of quinone exposure treatments (On average, they were 8.10% more likely to disperse from the 1^st^ patch of the linear array than females; Fig S4)

**3. SUPPLEMENTARY FIGURES**

**Figure S1.** A brief experimental outline of two-way-choice of scavenging between normal vs infected carcasses. 12-day-old mated females, in a group 3, were simultaneously provided access to normal vs infected carcasses (3 carcasses of each type, impaled with an insect pin) and assayed for the difference in scavenging activity (n=13 replicate groups/carcass type).

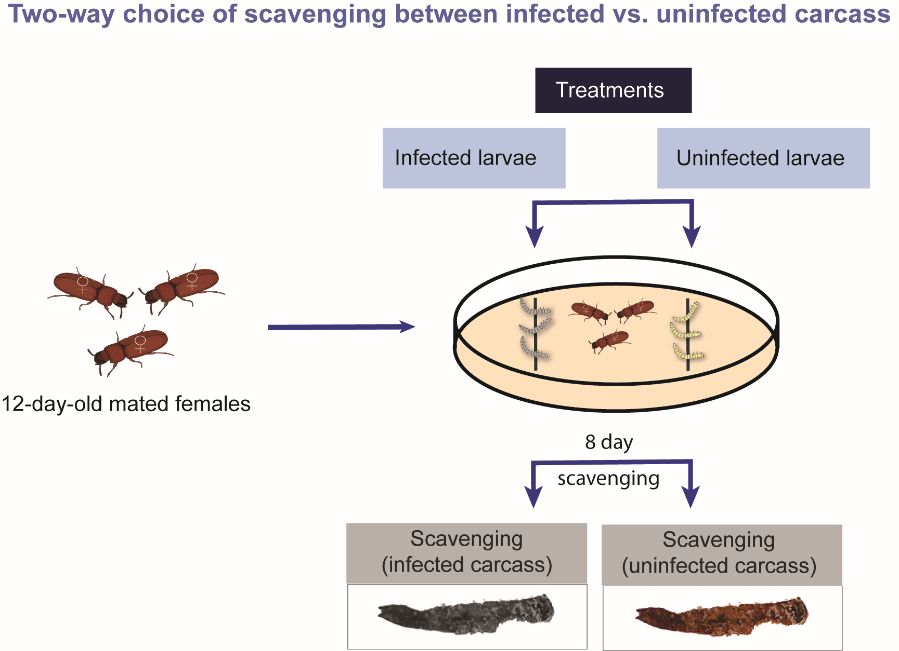

**Figure S2. Quinone production of individual females, estimated by the zone of inhibition assay** (Khan *et al.* 2015)**, strongly correlates with the amount of individual stink gland quinone components, quantified HPLC methods** (Khan *et al.* 2018) (A) Quantitation of total quinone production by females from MB vs FB groups**,** measured as the mean diameter (SE) of the zone of inhibition of bacterial growth on agar plates produced by individual cold-shocked females embedded in the soft agar plate (unpublished data, I Khan); (B-C) Data from (Khan *et al.* 2018), reanalysed to show the amount of (B) methyl benzoquinone (MBQ) and (C) ethyl benzoquinone (EBQ) in dissected stink glands of females from MB vs FB groups, measured by HPLC methods. P values in panels show the significant effects of sex-ratio.

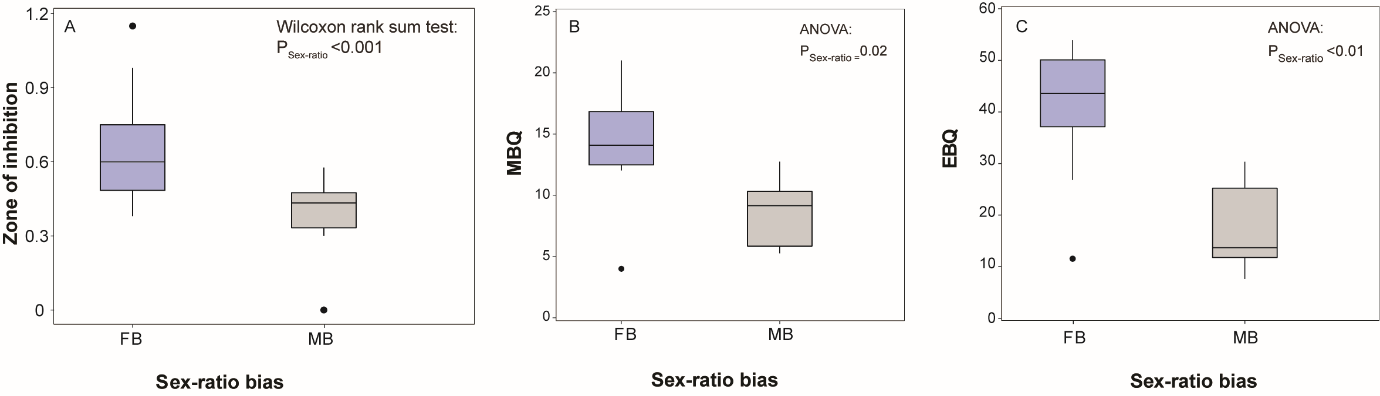

**Figure S3**: (A) Decumulative probability distributions for dispersal tendencies of males and females across sex-ratio groups (Male-biased: MB vs Female-biased: FB) (n=10 replicate arrays/sex-ratio bias). The X-axis represents distance dispersed by the individuals (in terms of replicate patch number) from the patch of origin (i.e. x= 0; where the beetles were initially released). Y-axis denotes the probability of further movement away than the distance already dispersed along the X-axis (e.g. while y-value at x = 0 denotes the probability of dispersal from the first patch, the y-value at x = 1 is the probability of dispersal at least to the third patch). P-value represents the two-way interaction between sex-ratio bias and sex, derived from the ordinal logistic model best fitted with logit linked function; (B) Dispersal kernels across sexes. Dotted lines represent the proportion of beetles found in different patches of individual linear arrays across sexes; whereas bold lines represent their means across all linear arrays. P-value represents results from the ordinal logistic model best fitted with logit link function; Average (C) Mean, (D) Standard deviation and (E) Maximum dispersal of individuals across sexes and sex-ratio groups from the simulation. Y-axis represents the dispersal distance away from the original patch (i.e. patch 1). We analysed each kernel parameter using a linear mixed effects model. Each panel shows points that averaged kernel parameters across 1000 runs, with 95% confidence intervals. P-values in C-E represent the average significance of the main effects as well as their interaction. SR= Sex-ratio groups; S= Sex.

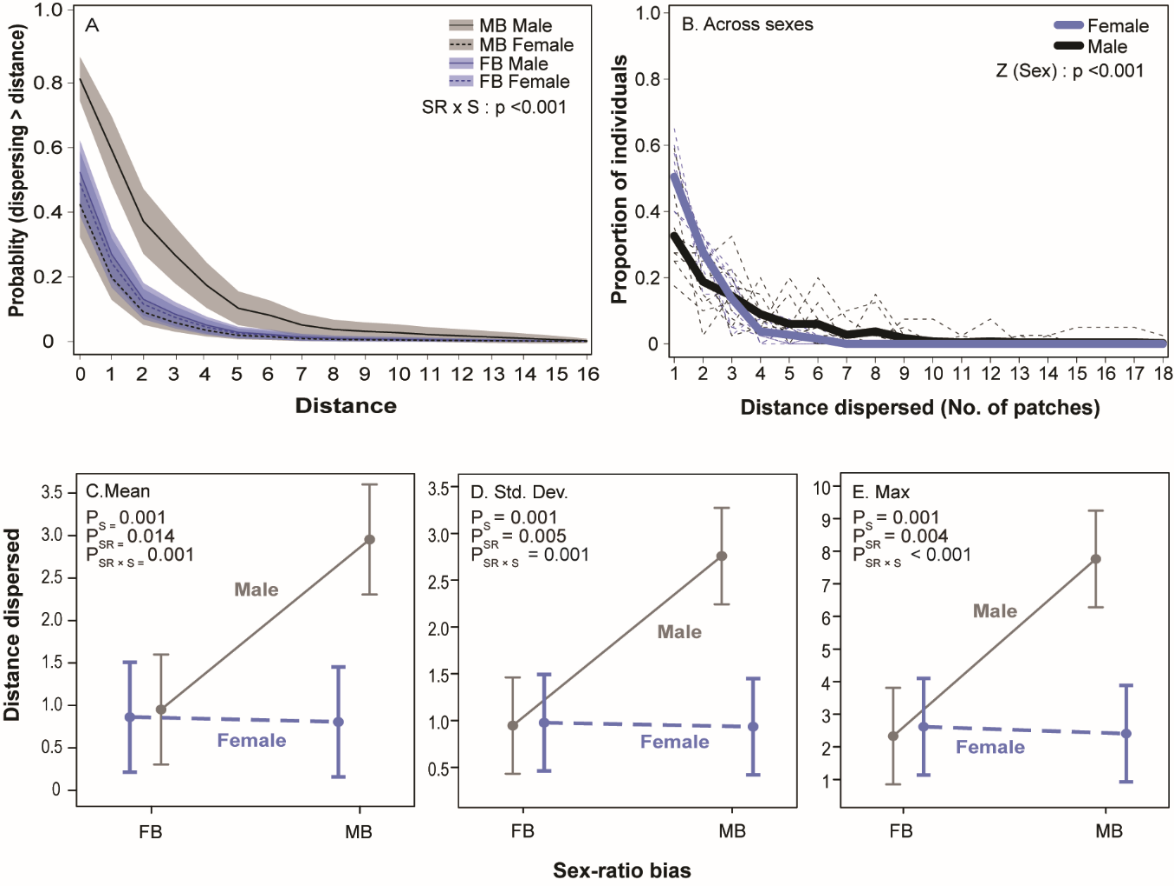

**Figure S4**. (A–D) Dispersal kernels for individuals from different treatments (e.g. MBQ exposure vs control) and sexes (male vs female). Comparison across (A) treatments (B) sexes; (C) females and (D) males separately across treatments (n=9 replicate arrays/treatment). Dotted lines represent the proportion of beetles found in different patches of individual linear arrays across sexes and quinone treatments; whereas bold lines represent their means across all linear arrays. P-values represent results for each comparison derived from the ordinal logistic model best fitted with logit link function; (E) Decumulative probability distributions of dispersal tendencies of males and females as a function of quinone exposure. Axes have been described in Fig. S3. We analysed the data using an ordinal logistic model best fitted with logit linked function. P-value represents the two-way interaction between quinone exposure and sex. S = Sex; T = Treatment.

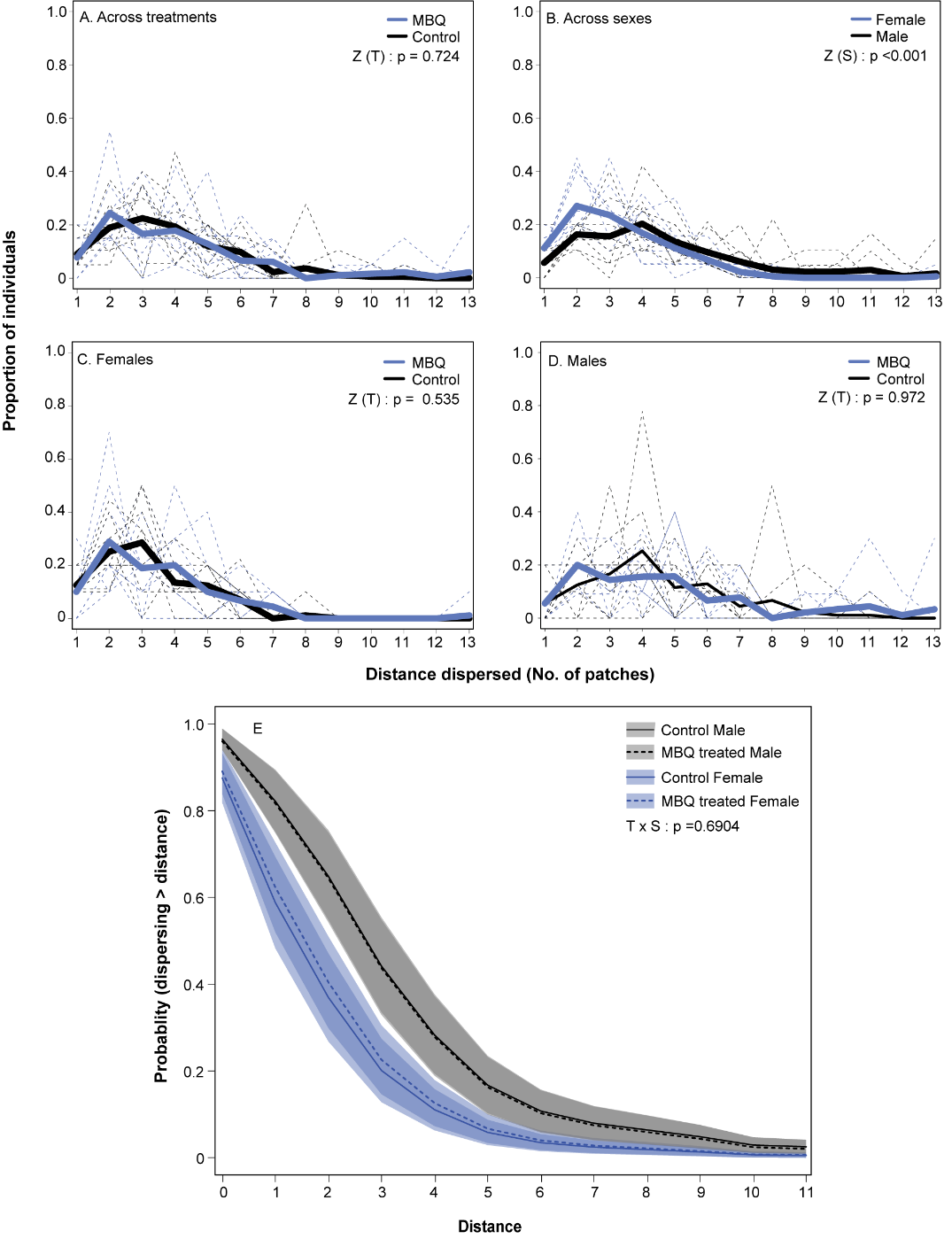

**4. SUPPLEMENTARY TABLES**

**Table S1: Analysis of scavenging and its impact on fecundity and stink gland production.** (A) Summary of ANCOVA for fecundity as a function of sex-ratio bias and available natal resource. (B) Summary of a generalised linear model for group scavenging as a function of sex-ratio bias and carcass type (Data fitted to a quasibinomial distribution) [Model specification: Scavenging activity ~ Sex-ratio $\times$ Sex]. (C) Summary of a generalised linear model for per capita scavenging activity as a function of sex-ratio bias and sex (Data fitted to a quasibinomial distribution) [Model specification: Scavenging activity ~ Sex-ratio $\times$ Carcass type]. (D) Summary of a linear mixed-effects model for fecundity as a function of carcass type and sex-ratio bias, using replicate experiment as a random factor [Model specification: Fecundity ~ Sex-ratio $\times$ Carcass type + (1| Replicate experiment)]. (E) Summary of ANOVA for beetle quinone production estimated as a normalised zone of inhibition as a function of sex-ratio bias and carcass type. (F) Summary of paired t-test for scavenging preference between infected vs normal carcasses.

| **Experiment, effect** | df | SS | F value | P |
| --- | --- | --- | --- | --- |
| (A) *Fecundity with different amounts of natal resource (Fig 2A)* |  |  |  |  |
| Sex-ratio | 1 | 505.704 | 31.007 | **<0.001** |
| Resource | 1 | 144.611 | 8.866 | **0.04** |
| Sex-ratio $\times$ Resource | 1 | 3.861 | 0.236 | 0.628 |
| Residuals | 60 | 978.547 |  |  |
| **Experiment, effect** |  | **df** | **χ^2^** | **P** |
| (B) *Group scavenging (Fig 3A)* |  |  |  |  |
| Sex-ratio |  | 2 | 193.736 | **< 0.001** |
| Carcass type |  | 1 | 84.478 | **< 0.001** |
| Sex-ratio × Carcass type |  | 2 | 55.23 | **< 0.001** |
| (C) *Per capita scavenging (Fig 3B)* |  |  |  |  |
| Sex-ratio |  | 1 | 6.218 | **0.01** |
| Sex |  | 1 | 42.344 | **<0.001** |
| Sex-ratio × Sex |  | 1 | 3.567 | **0.059** |
| (D) *Effect of scavenging on fecundity (Fig 3C)* |  |  |  |  |
| Sex-ratio |  | 1 | 24..23 | **<0.001** |
| Carcass type |  | 2 | 44.41 | **<0.001** |
| Sex-ratio × Carcass type |  | 2 | 25.96 | **<0.001** |
| **Experiment, effect** | **df** | **SS** | **F-value** | **P** |
| (E) *Effect of scavenging on quinone production (Fig 3D)* |  |  |  |  |
| Sex-ratio | 1 | 0.3921 | 45.725 | **<0.001** |
| Carcass type | 2 | 0.1672 | 9.746 | **<0.001** |
| Sex-ratio × Carcass type | 2 | 0.0713 | 4.155 | **0.02** |
| Residuals | 87 | 0.746 |  |  |
| **Experiment, effect** |  | **df** | **t-statistic** | **P** |
| (F) *Choice between infected vs uninfected carcass (Fig* 3E) |  |  |  |  |
| Carcass type |  | 12 | 4.1262 | **0.001** |

**Table S2: Analysis of dispersal behaviour across sexes and sex-ratio group.** (A) Results of the ordinal logistic regression analysis for the effects of sex-ratio bias and sex on the dispersal kernel of beetles across the linear arrays. Kernel parameters with significant effects have been shown separately. μ = mean, ∨ = maximum, σ = standard deviation. (B) Summary of average χ^2^ and p-values from linear mixed models showing the effects of sex-ratio and sex, and their interaction on 5 dispersal kernel parameters (μ, ∨, σ, γ = skew, and κ = kurtosis) [Model: Distance dispersed ~ Sex-ratio bias × Sex + (1|replicate array)]. The results reported above are averaged across the 1000 iterations of simulated data for Type II SS ANOVA for each kernel parameters (*car* package, Fox & Weisberg 2019)).

|  | **A. Effects** | | | **LR stat** | | | | **P** | | | | **Parameters** | | |
| --- | --- | --- | --- | --- | --- | --- | --- | --- | --- | --- | --- | --- | --- | --- |
|  | Sex (S) | | | 23.173 | | | | **<0.001** | | | | μνσ | | |
|  | SR $\times$ S | | | 18.749 | | | | **<0.001** | | | | μνσ | | |
|  | Sex-ratio bias (SR) | | | 8.692 | | | | **0.003** | | | | μνσ | | |
| **B. Effects** | | **Mean** | | | **SD** | | **Skewness** | | | **Kurtosis** | | | **Maximum** | |
|  |  | χ^2^ | P | | χ^2^ | P | χ^2^ | | P | χ^2^ | P | | χ^2^ | P |
| SR | | 8.285 | **0.014** | | 11.475 | **0.005** | 1.468 | | 0.4 | 1.059 | 0.476 | | 11.792 | **0.004** |
| S | | 18.926 | **0.001** | | 15.382 | **0.001** | 2.886 | | 0.275 | 1.355 | 0.435 | | 15.108 | **0.001** |
| SR$\times$ S | | 16.46 | **0.001** | | 16.461 | **0.001** | 1.184 | | 0.474 | 1.418 | 0.453 | | 18.629 | **<0.001** |

**Table S3: Analysis of dispersal behaviour across sexes and quinone exposure treatments.** Results of the ordinal logistic regression analysis for the effects of quinone exposure and sex on the dispersal kernel of beetles across the linear arrays [Model: Distance dispersed ~ Quinone exposure treatment × Sex + (1|replicate array)].

| **Effects** | **LR stat** | **P** |
| --- | --- | --- |
| Treatment | 0.1606 | 0.688 |
| Sex | 29.698 | **<0.001** |
| Treatment $\times$ Sex | 0.1586 | 0.69 |

**Table S4: Analysis of scavenging as a protein source**. (A) Summary of generalised linear model for scavenging activity as a function of sex-ratio bias and protein source (Data fitted to quasibinomial distribution) [Model specification: Scavenging rate ~ Sex-ratio $\times$ Protein source]. (B) Summary of generalised linear model for fecundity as a function of sex-ratio bias and protein source (Data fitted to Poisson distribution) [Model specification: Fecundity ~ Sex-ratio $\times$ Protein source].

| **Experiment, effect** |  | df | $\chi$^2^ | P |
| --- | --- | --- | --- | --- |
| (A) *Scavenging (Fig 4A)* |  |  |  |  |
| Sex-ratio |  | 1 | 84.889 | **<0.001** |
| Protein source |  | 1 | 241.268 | **<0.001** |
| Sex-ratio $\times$ Protein source |  | 1 | 4.449 | **0.04** |
| (B) *Offspring per female*   *(Fig 4B)* |  |  |  |  |
| Sex-ratio |  | 1 | 12.97 | **<0.001** |
| Protein source |  | 3 | 24.73 | **<0.001** |
| Sex-ratio $\times$ Protein source |  | 3 | 6.64 | 0.08 |

**Table S5: Analysis of stink gland contents on scavenging activity and fecundity.** (A) Summary of generalised linear model for scavenging activity as a function of abdominal stink gland extract or individual stink gland components (i.e. MBQ and PD) (Data fitted to quasibinomial distribution) [Model specification: Scavenging activity~abdominal stink gland extract or MBQ or PD]. (B) Summary of ANOVA for fecundity per female as a function of abdominal stink gland extract or individual stink gland components (with or without the access to carcass). Effects of stink gland extract, MBQ and PD were analysed separately.

| **Experiment, effect** |  | df | $\chi$^2^ | P |
| --- | --- | --- | --- | --- |
| *A. Fraction of larvae remaining as a function of stink gland extract or individual stink gland components (Fig 5A, C)* |  |  |  |  |
| Stink gland extract |  | 1 | 5.222 | **0.02** |
| MBQ |  | 1 | 11.766 | **<0.001** |
| PD |  | 1 | 9.91 | **0.002** |
| **Experiment, effect** | df | SS | F value | P |
| *B. Fecundity as a function of stink gland extract or individual stink gland components (Fig 5B, D)* |  |  |  |  |
| Stink gland extract | 3 | 385.8 | 9.37 | **<0.001** |
| Residuals | 72 | 988.1 |  |  |
| MBQ | 2 | 208.2 | 9.12 | **0.001** |
| Residuals | 57 | 650.7 |  |  |
| PD | 2 | 102.7 | 4.298 | **0.02** |
| Residuals | 57 | 681 |  |  |

**5. SUPPLEMENTARY REFERENCES**

Bates, D., Mächler, M., Bolker, B.M. & Walker, S.C. (2015). Fitting linear mixed-effects models using lme4. *J. Stat. Softw.*, 67.

Christensen, R.H.B. (2019). ordinal—Regression models for ordinal Data. *R package* *version* 2019.12-10., 54.

Endriss, S.B., Vahsen, M.L., Bitume, E. V., Grey Monroe, J., Turner, K.G., Norton, A.P., *et al.* (2019). The importance of growing up: juvenile environment influences dispersal of individuals and their neighbours. *Ecol. Lett.*, 22, 45–55.

Fox, J. & Weisberg, S. (2019). *An R companion to applied regression,* Third edition. Sage, Thousand Oaks, CA.

Khan, I., Prakash, A. & Agashe, D. (2015). Immunosenescence and the ability to survive bacterial infection in the red flour beetle *Tribolium castaneum*. *J. Anim. Ecol.*, 85, 291–301.

Khan, I., Prakash, A., Issar, S., Umarani, M., Sasidharan, R., Masagalli, J.N., *et al.* (2018). Female density-dependent chemical warfare underlies fitness effects of group sex ratio in flour beetles. *Am. Nat.*, 191, 306–317.

Lenth, R., Singmann, H., Love, J., Buerkner, P. & Herve, M. (2020). Package ‘emmeans.’ *R Package version 1.4.6.*, 34, 216–221.

Lenth, R. V. (2016). Least-squares means: The R package lsmeans. *J. Stat. Softw.*, 69.

Li, J., Lehmann, S., Weißbecker, B., Ojeda Naharros, I., Schütz, S., Joop, G., *et al.* (2013). Odoriferous defensive stink gland transcriptome to identify novel genes necessary for quinone synthesis in the red flour beetle, *Tribolium castaneum*. *PLoS Genet.*, 9, e1003596.

Prendeville, H.R. & Stevens, L. (2002). Microbe inhibition by *Tribolium* flour beetles varies with beetle species, strain, sex, and microbe group. *J. Chem. Ecol.*, 28, 1183–90.

Unruh, L.M., Xu, R. & Kramer, K.J. (1998). Benzoquinone levels as a function of age and gender of the red flour beetle , *Tribolium castaneum*, *Insect Biochem. Mol. Biol.,* 28, 969–977.
